# A transferrable and interpretable multiple instance learning model for microsatellite instability prediction based on histopathology images

**DOI:** 10.1101/2020.02.29.971150

**Authors:** Rui Cao, Fan Yang, Si-Cong Ma, Li Liu, Yan Li, De-Hua Wu, Yu Zhao, Tong-Xin Wang, Wei-Jia Lu, Wei-Jing Cai, Hong-Bo Zhu, Xue-Jun Guo, Yu-Wen Lu, Jun-Jie Kuang, Wen-Jing Huan, Wei-Min Tang, Junzhou Huang, Jianhua Yao, Zhong-Yi Dong

## Abstract

**Background:** Microsatellite instability (MSI) is a negative prognostic factor for colorectal cancer (CRC) and can be used as a predictor of success for immunotherapy in pan-cancer. However, current MSI identification methods are not available for all patients. We propose an ensemble multiple instance learning (MIL)-based deep learning model to predict MSI status directly from histopathology images.

**Design:** Two cohorts of patients were collected, including 429 from The Cancer Genome Atlas (TCGA-COAD) and 785 from a self-collected Asian data set (Asian-CRC). The initial model was developed and validated in TCGA-COAD, and then generalized in Asian-CRC through transfer learning. The pathological signatures extracted from the model are associated with genotypes for model interpretation.

**Results:** A model called Ensembled Patch Likelihood Aggregation (EPLA) was developed in the TCGA-COAD training set based on two consecutive stages: patch-level prediction and WSI-level prediction. The EPLA model achieved an area-under-the -curve (AUC) of 0.8848 in the TCGA-COAD test set, which outperformed the state-of-the-art approach, and an AUC of 0.8504 in the Asian-CRC after transfer learning. Furthermore, the five pathological imaging signatures identified using the model are associated with genomic and transcriptomic profiles, which makes the MIL model interpretable. Results show that our model recognizes pathological signatures related to mutation burden, DNA repair pathways, and immunity.

**Conclusion:** Our MIL-based deep learning model can effectively predict MSI from histopathology images and are transferable to a new patient cohort. The interpretability of our model by association with genomic and transcriptomic biomarkers lays the foundation for prospective clinical research.

## INTRODUCTION

Microsatellite instability (MSI) is a hypermutator phenotype that occurs in tumors with impaired DNA mismatch repair (MMR)[1], which is reported as a hallmark of hereditary Lynch syndrome (LS)-associated cancers[2] and observed in about 15% of colorectal cancer (CRC)[3]. High levels of MSI (MSI-H) have been found to be a favorable prognostic factor but a negative predictor for adjuvant chemotherapy in stage II CRC[4]. More importantly, recent studies have demonstrated MSI-H/MMR deficiency is correlated to an increased neoantigen burden, which sensitizes the tumor to immune checkpoint blockade (ICB) treatment[5]. Further investigations suggest that the benefit of ICB treatment for patients with MSI-H is not limited to specific tumor types but to all solid tumors[6], which establishes the crucial role of MSI in predicting the efficacy of immunotherapy for advanced solid tumors, especially CRC.

MSI or MMR-deficiency (dMMR) testing has traditionally been performed in patients with CRC and endometrial cancer (EC) to screen for LS–associated cancer predisposition[7]. Recently, with the U.S. Food and Drug Administration (FDA) designation of MSI/dMMR as a favorable predictor of anti-PD-1 immunotherapy[8] (a subset of ICB treatment), the clinical demand for MSI/dMMR testing has increased dramatically. However, in clinical practice, not every patient is tested for MSI, especially in those cancers with lower occurrences of MSI or in patients in developing countries, because it requires additional genetic or immunohistochemical tests which are costly and time-consuming. Additionally, various existing MSI testing methods show different sensitivities and specificities, leading to the disunity of results[9, 10]. Therefore, there are both opportunities and challenges that lie ahead in developing an MSI testing method that is available for all cancer patients.

The emergence of computational pathology had provided an opportunity for the detection of MSI because pathology slides are produced for almost every patient diagnosed with cancer; these slides can be digitized into whole slide images (WSIs)[11]. WSI can not only reveal the tissue spatial arrangement of tumor cells at low magnification, but also the cell structure at high magnification[12]. Furthermore, histopathology images also show the immunologic microenvironment of tumors[13]. Thus, the cell level phenotypes expressed in WSI are the direct reflection of genotypes such as MSI at the molecular scale. With the continuous penetration of artificial intelligence (AI) into the field of medical imaging, researchers have sought solutions based on deep learning, a subset of AI, in a wide range of medical problems, such as prediction of gene mutation[14] and tumor-infiltrating lymphocytes[12]. Whereas traditional machine learning depends largely on the rationality of human-selected features[15], deep learning can learn features from the data, which makes it possible for researchers to discover uncharted findings[16, 17]. Previous studies have suggested that deep learning can discover regions that contribute to MSI status with special pathomorphological characteristics[18], but the applicability of the model in the Asian population remains in question because of the great variation in demographics and data preparation. The inability to interpret the extracted features and the predictions made by the model is considered to be one of the major issues that limits the acceptance of AI models in medicine.

In this study, we developed a multiple-instance-learning (MIL)-based deep learning model to predict MSI status from histopathology images. The model, which we have termed Ensemble Patch Likelihood Aggregation (EPLA), combines both deep learning and traditional machine learning techniques. The proposed MIL model was trained using the TCGA-CORD data set, and then transfer learning was implemented to fine tune the model using the Asian-CRC data set, which enhanced the generalizability of this model. More importantly, we also demonstrate the interpretability of the model by identifying the critical MIL model generated pathological signatures and linking them with MSI genomic and transcriptomic profiles.

## MATERIALS AND METHODS

### Patient cohorts and dataset partition

In this study, whole slide images (WSI) of two large cohorts were collected. An MSI label was assigned to each WSI based on the patient’s microsatellite measurement, where samples with MSI sensor scores less than 10 were denoted as microsatellite stable tumors (MSS) and those greater than or equal to 10 were denoted as microsatellite instable (MSI)[19]. The first cohort (TCGA-COAD), retrieved from The Cancer Genome Atlas, comprised 429 frozen tissue slides diagnosed as colon adenocarcinoma and were downloaded from the Digital Slide Archive (https://cancer.digitalslidearchive.org/); 358 cases were labeled as MSS and 71 cases were labeled as MSI. The second cohort (Asian-CRC), collected from Tongshu Biotechnology Co., Ltd, consisted of 785 formalin-fixed paraffin-embedded (FFPE) sections diagnosed as CRC, which were provided from three medical centers in China. Patients in the Asian-CRC group were analyzed by an MSI detection kit (Shanghai Tongshu Biotechnology Co., Ltd.), and 164 cases were identified as MSI-high; the balance cases (621 cases) were identified as MSI-low or MSS. In order to use these histopathology images, patient consents were obtained, and this study was approved by the Institutional Ethical Review Boards of Nanfang Hospital. The details of the two cohorts are summarized in Table S1.

The TCGA-COAD cohort was split into separate training and test sets at a 7:3 ratio. In order to maintain the same ratio of positive to negative samples in the training set and test set, a data splitting procedure was conducted as follows: (1) all WSIs in the dataset were divided into two groups, i.e, the WSI group with the MSI label, and the group with the MSS label; (2) 70% images in the MSI group and 70% images in the MSS group were randomly selected to compose the training set; and (3), the remaining images made up the test set. The training set was used for hyper-parameter tuning based on cross-validation, whereas the test set was used for the evaluation of generalization performance.

### ROI delineation, tiling, and data preprocessing

All WSIs were digitalized at 20× magnification with a predefined pixel size (=0.5μm). In order to reduce the influence of unrelated areas and alleviate the workload of the classification method, regions of carcinoma (ROI) on WSIs were manually annotated by expert pathologists, according to the following rules: (1) the tumor cells should occupy more than 80% of an ROI, i.e., the interstitial component is less than 20%; and (2), obvious interfering factors, including creases, bleeding, necrosis and blurred areas, should be excluded. The annotation was performed using Aperio ImageScope (Aperio Technologies, Inc.).

Given the extremely large image size (typically 100,000 × 50,000 pixels) of a WSI, the WSIs were subsequently tiled into 512×512 patches. Only patches having a greater than 80% overlap with the carcinoma ROI were used for the following analysis. The number of patches per WSI in TCGA-COAD ranged from 22 to 2357 (average 224), whereas the Asian-CRC ranged from 5 to 3718 (average 338) (Table S1).

Data augmentation and normalization were applied for training patches, whereas only normalization was employed for test patches. Data augmentations used in our work included random horizontal flipping and random affine transformation of the patches (keeping the center invariant). The random affine transformation had the following processes: (1) a random rotation ranging from −90 to +90 degrees; and (2), a random translation with the horizontal shift and the vertical shift range from −8% to +8% of patch width and height respectively. Finally, the augmented patches were center cropped to 224 pixels × 224 pixels, following a z-score normalization on RGB channels.

### MIL-based Deep Learning pipeline

Our MIL-based deep learning pipeline is illustrated in Figure 1 (‘Processing a WSI in Two Stages’). Figure 1 presents two predictions: patch-level and WSI-level. Due to the large image size, the WSI was first divided into small patches, and then the patch likelihoods were aggregated in an ensemble classifier to obtain the WSI-level prediction. Therefore, our method is termed Ensemble Patch Likelihood Aggregation (EPLA).

**Figure 1.**
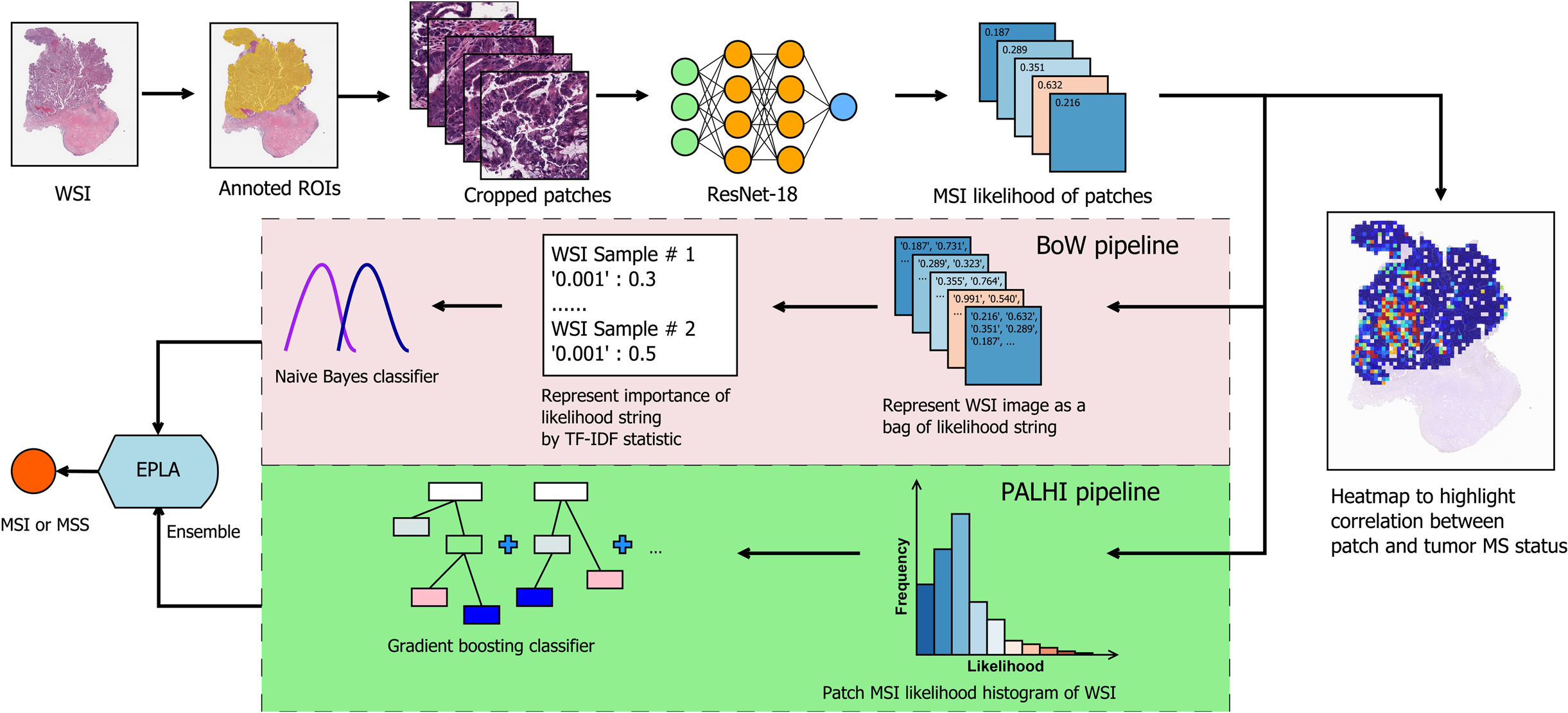
Overview of the Ensemble Patch Likelihood Aggregation (EPLA) model. A WSI of each patient was obtained and annotated to highlight the regions of carcinoma (ROIs). Then, patches were tiled from ROIs, and the MSI likelihood of each patch was predicted by ResNet-18, during which a heat map was shown to visualize the patch-level prediction stage. Next, PALHI and BoW pipelines integrated the multiple patch-level MSI likelihoods into a WSI-level MSI prediction value, respectively. Finally, ensemble learning combined the results of the two pipelines and made the final prediction of the MSI status.

During the patch-level prediction, a residual convolutional neural network (ResNet-18) was trained to compute the patch likelihood in a MIL paradigm where the patches were assigned with the WSI’s label. Binary cross-entropy (BCE) loss was utilized to optimize the network using a mini-batch gradient descent method.

We used two independent MIL methods to aggregate the patch likelihoods: Patch Likelihood Histogram (PALHI) pipeline and Bag of Words (BoW) pipeline, which were inspired by the histogram-based method[20] and the vocabulary-based method[21], respectively. In PALHI, a histogram of the occurrence of the patch likelihood was applied to represent the WSI, whereas in BoW, each patch was mapped to a TF-IDF floating-point variable, and a TF-IDF feature vector was computed to represent the WSI.

Traditional machine learning classifiers were then further trained using these feature vectors to predict the MSI status for each WSI. Here, Extreme Gradient Boosting (xgboost), a kind of gradient boosted decision tree, was employed in the PALHI pipeline. Naïve Bayes (NB) was used in the BoW pipeline. During the training of the WSI-level classifier, the hyperparameters were determined based on the cross-validation on the training set, using WSI-level ROCAUC (ROCAUC) as the performance metric. The PALHI and BoW classifiers were then ensembled to obtain the final prediction[22].

### Transfer learning

Transfer learning applies the knowledge learned in one field to different but related fields. The initial parameters of the model were trained using the public data of TCGA-COAD, and the transfer learning was implemented using the Asian-CRC data. The transfer learning was conducted by reusing the model weights in the patch-level discriminators and then fine-tuning the weights using a small amount of labeled Asian-CRC data. In addition, we gradually added more Asian-CRC data for model finetuning to explore the impact on model performance, following the work by Yosinski et al[23].

### Model interpretability through molecular correlation analysis of pathological signatures

We explored our model interpretability by examining the pathological signatures and their correlation with MSI-related genome types.

#### Identification of pathological signatures of importance

The occurrence histograms in the PALHI and the TF-IDF feature vector in BoW were the pathological signatures generated by our model. The importance of each signature was measured by its contribution weight to the final WSI-level prediction. Wilcoxon Rank Sum tests were performed to evaluate the discriminating power of each pathological signatures. The top signatures were then sent for genomic and transcriptomic correlation analysis.

#### Genomic and transcriptomic data acquisition

The DNA mutation profile and the mRNA expression profile of TCGA-COAD were retrieved from cBioPortal (http://www.cbioportal.org/)[24]. The RNA-seq data was normalized using RSEM[25]. The synonymous mutations were excluded from the following correlation analysis. For a particular gene set, as long as there was a non-synonymous mutation in any of its gene members, it would be defined as deficient.

#### Genomic correlation analysis

The relationship between MSI and some mutation indexes has been reported in previous literature, including INDEL and TMB[26]. INDEL mutations refer to a variant type caused by sequence insertion (INS) or deletion (DEL) and can be calculated as the frequency of DEL and INS mutations. As the mutation data was profiled by the whole exome sequencing, TMB was defined and calculated as the total number of somatic nonsynonymous mutations of the entire genome[27]. To explore the relationship between the pathological signatures and these known genomic biomarkers, they were first normalized to a range of 0 to 1 and then visualized in a heat map using an R package pheatmap, during which unsupervised clustering was applied using Ward’s minimum variance method.

#### Transcriptomic correlation analysis

WGCNA is a bioinformatics method based on expression data and is typically used to identify gene modules with highly synergistic changes[28]. We first constructed a co-expression network for the mRNA expression profile using the R package WGCNA, during which the soft threshold for the network was set to the recommended value selected by the function pickSoftThreshold (Figure S1). Setting the minimum module size to 100 and other parameters to default, we identified 24 transcriptomic modules (Figure S2). The biological functions of the modules were annotated by the Gene Ontology (GO) over-representation test using the R package clusterProfiler[29], during which the Benjamini-Hochberg method was used to adjust *P* value. Only those GO terms with adjusted *P* values lower than 0.05 were considered significantly enriched in a particular module. After that, we calculated Spearman’s rank correlation coefficients for each pair of modules and pathological signatures to recognize the modules in interest.

### Statistical analysis

The ROC curves were drawn using pROC and ggplot2 in R (version 3.6.1). The area under the ROC curve was calculated in pROC. The significance of AUC differences was tested using the Wald test statistic. The optimal cutoff points of the ROC curves were estimated using the Youden Index. The Wilcoxon Rank Sum test was used to compare two paired groups and visualized as a boxplot using R package ggpubr. Spearman’s rank correlation coefficients were used for correlation analysis.

## RESULTS

### Development of Ensemble Patch Likelihood Aggregation (EPLA)

Our model consists of two consecutive stages: patch-level prediction and WSI-level prediction (Figure 1). Briefly, a WSI was annotated to delineate the region of carcinoma (ROI). The ROI was tiled into patches, which were subsequently fed to a residual convolutional neural network (ResNet-18) to obtain the patch-level MSI prediction. Then, we trained two independent MIL pipelines to integrate multiple patch-level predictions into an MSI value at the WSI level: the PAtch Likelihood HIstogram (PALHI) pipeline and the Bag of Words (BoW) pipeline. To obtain the optimal convex combination of the two MIL methods, we employed ensemble learning to eventually obtain the predicted MSI status of the patient (Figure 1).

### Performance of EPLA

The performance of the EPLA model was measured in the TCGA-COAD test set. Two representative heat maps, MSI and MSS respectively, are shown in Figure 2A. These heat maps provide the patch-level prediction. The EPLA model achieved an AUC of 0.8848 at the WSI level (Figure 2B) and outperformed the state-of-the-art deeplearning based majority voting method (denoted as DL-based MV) in Kather’s study[18], which trains a ResNet for patch-level predictions and then takes the majority of these predictions as the final MSI status of the patient (Figure 2D). To directly compare our method with the DL-based MV in the same test set, we implemented DL-based MV method as a variant of PALHI, by downsizing the histogram in PALHI to only two bins (0-0.5 and 0.5-1). Consistent with the results in Kather’s study, the DL-based MV method achieved similar performance with an AUC of 0.8457 in the TCGA test set (Figures 2B and 2D).

**Figure 2.**
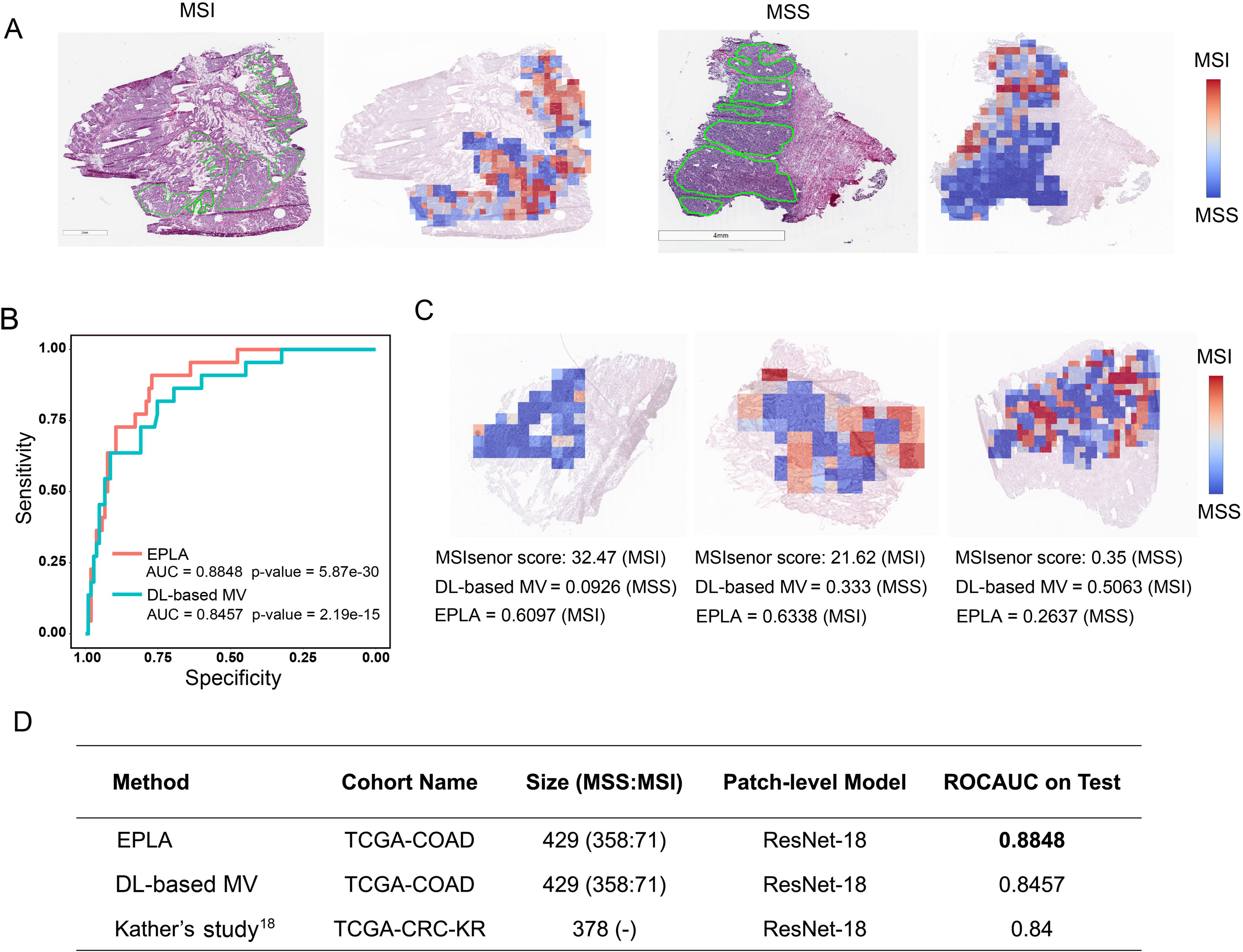
Validation of the EPLA and comparison with DL-based MV in the TCGA cohort. (A) Representative heat maps of MSI and MSS cases at the patch-level prediction stage. Color bars show the MSI likelihood of each patch. (B) Receiver operating characteristic (ROC) curves of EPLA and DL-based MV. The *P* value is calculated by the Wald test. (C) A heat map of representative discrepant cases between EPLA and DL-based MV. (D) Summary of EPLA and DL-based MV. DL-based MV is re-implemented from a voting-based model in Ref. 18. The last line of the table summarizes the performance of the original DL-based MV model.^18^

We further compared the performance of components of the EPLA ensemble (i.e., PALHI and BoW) to that of DL-based MV (Table S2). We found that BoW achieved higher specificity (89.5% vs 75.2%) and PALHI was superior in terms of sensitivity (86.4% vs 81.8%). The ensembled EPLA classifier combines the advantage of both models, which obtained both superior specificity and sensitivity compared to the DL-based MV (Table S2). Representative heat maps of the discrepant cases are shown in Figure 2C. These cases were correctly predicted by EPLA but mistakenly classified by DL-based MV.

### Validation of EPLA in an Asian-CRC cohort

We further measure the generalizability of our model in an Asian-CRC cohort. It is noteworthy that there exist great differences between the Asian-CRC cohort and the TCGA-COAD cohort, not only in race but also in the slide preparation techniques (Table S1). As a consequence, when directly tested in Asian-CRC, the EPLA model trained on TCGA-COAD only achieved an AUC of 0.6497 (Figure 3A). Considering the wide variations in medical practice, we applied transfer learning to generalize the EPLA model by fine-tuning ResNet-18 using 10% of cases from Asian-CRC, and thus achieved an AUC of 0.8504 in the remaining data set (Figures 3A and 3B).

**Figure 3.**
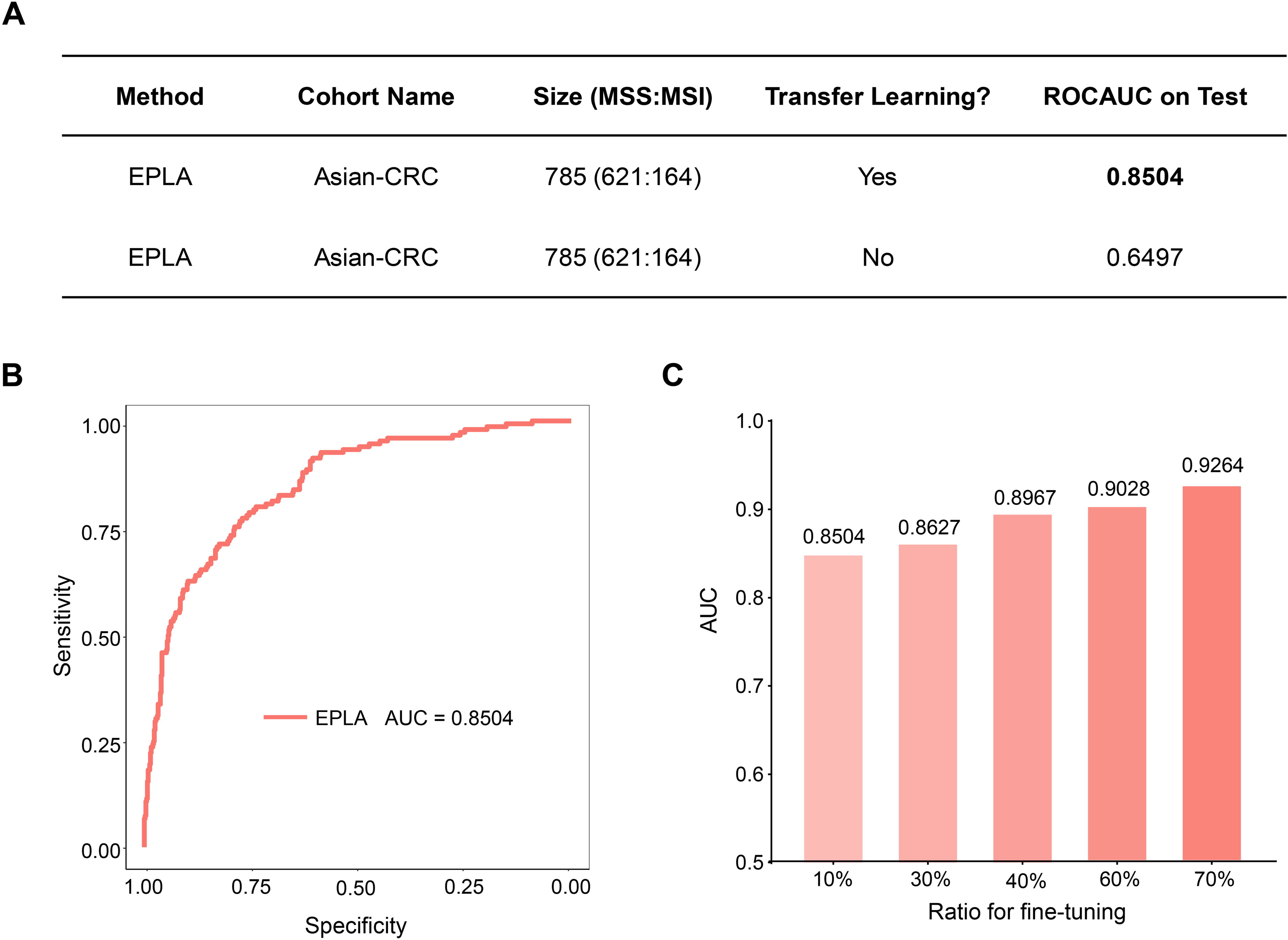
Generalization performance of the EPLA in an Asian cohort. (A) Summary of the performance of EPLA in Asian-CRC with or without transfer learning. When using transfer learning, 10% of cases from Asian-CRC were used for model fine-tuning. (B) The ROC curve of EPLA in the Asian-CRC using transfer learning with 10% of cases for model fine-tuning. (C) ROCAUCs of the model in Asian-CRC with increasing proportions of cases for model fine-tuning.

We subsequently evaluated the amount of data needed for model fine-tuning by increasing the proportion of cases from Asian-CRC for model training. The performance of the fine-tuned model steadily improved, resulting in 0.8627, 0.8967, 0.9028 and 0.9264 AUCs in the ratios of 30%, 40%, 60% and 70%, respectively, implying that transfer learning using a larger amount of data from a different cohort is the key to generalizing a model with a better performance. (Figure 3C).

### Top pathological signatures of the model

To gain insight into the MSI prediction mechanism of the model, we explored the contribution of the pathological signatures extracted from the EPLA model to the prediction of MSI in TCGA-COAD. The ranking of significance of the top ten pathological signatures is shown in Figure 4A. Given that the top five pathological signatures (FEA# 197, FEA# 198, FEA#001, FEA# 188 and FEA#200) were significantly more important than the others, they were selected for subsequent analysis. Among them, FEA#001 had a significantly higher value (P<0.0001) for patients in the MSS group, while the other four (FEA#188/197/198/200) had significantly higher values (P<0.0001) in the MSI group (Figure 4B).

**Figure 4.**
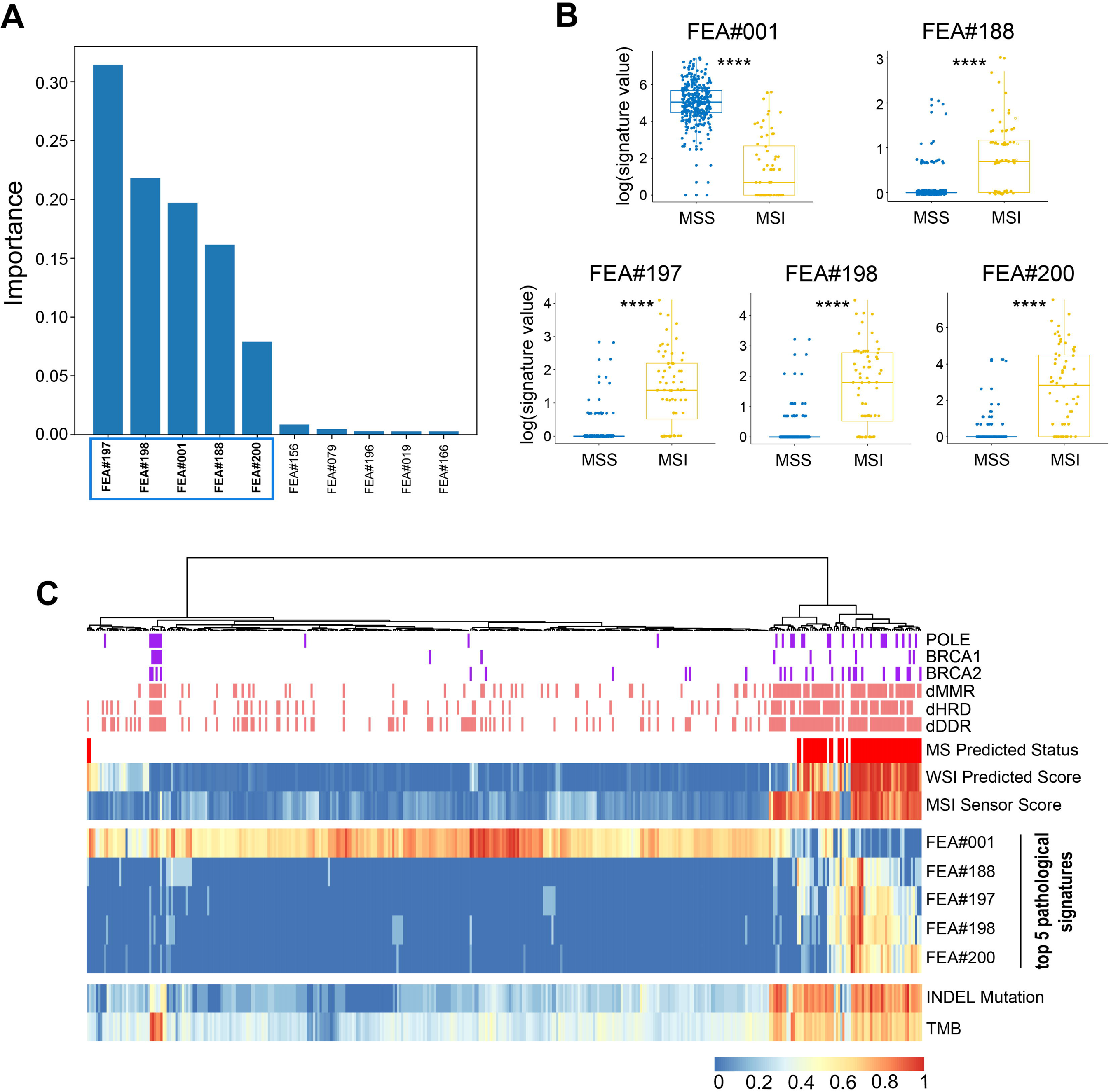
Identification and genomic correlation analysis of top pathological signatures. (A) Importance ranking of the top ten pathological signatures extracted from EPLA. (B) Boxplots of the five pathological signatures between MSI and MSS groups. Significance values: **** *P* < 0.0001. (C) Heat map with unsupervised clustering showing the correlation between genomic landscape and top pathological signatures in each patient. Each column corresponds to a patient in the TCGA-COAD cohort. All continuous variables are normalized to a range of 0 to 1. INDEL: insertion-deletion, TMB: tumor mutation burden, MMR: mismatch repair, DDR: DNA damage response and repair, and HRD: homologous recombination deficiency.

### Genomic association analysis

The output of the model can be interpreted from the perspective of the genome. Cluster analysis in figure 4C shows that patients with a high value of FEA#001 are mainly MSS with normal function in DNA repair-related pathways consisting of mismatch repair (MMR), DNA damage response and repair (DDR), and homologous recombination deficiency (HRD). On the contrary, patients with high levels of FEA#188/197/198/200 are mainly due to MSI with deficient DNA repair related pathways, namely deficient-MMR (dMMR), deficient-DDR (dDDR) and deficient-HRD (dHRD). In addition, mutations of several representative genes in these pathways, including POLE, BRCA1, and BRCA2, also demonstrate a consistent finding. Moreover, since recent evidence suggests MSI is significantly related to TMB, especially INDEL mutation load[26], we assessed the relation between the pathological signatures and these known biomarkers and found that high TMB and INDEL mutation load are often accompanied by low FEA#001 and high FEA#188/197/198/200 (Figure 4C).

### Transcriptomic association analysis

In transcriptomic association analysis of the top five pathological signatures, we applied weighted gene co-expression network analysis (WGCNA) and identified 24 modules (Figure 5A). Gene ontology (GO) enrichment analyses were performed to annotate the function of the modules (Table S3).

**Figure 5.**
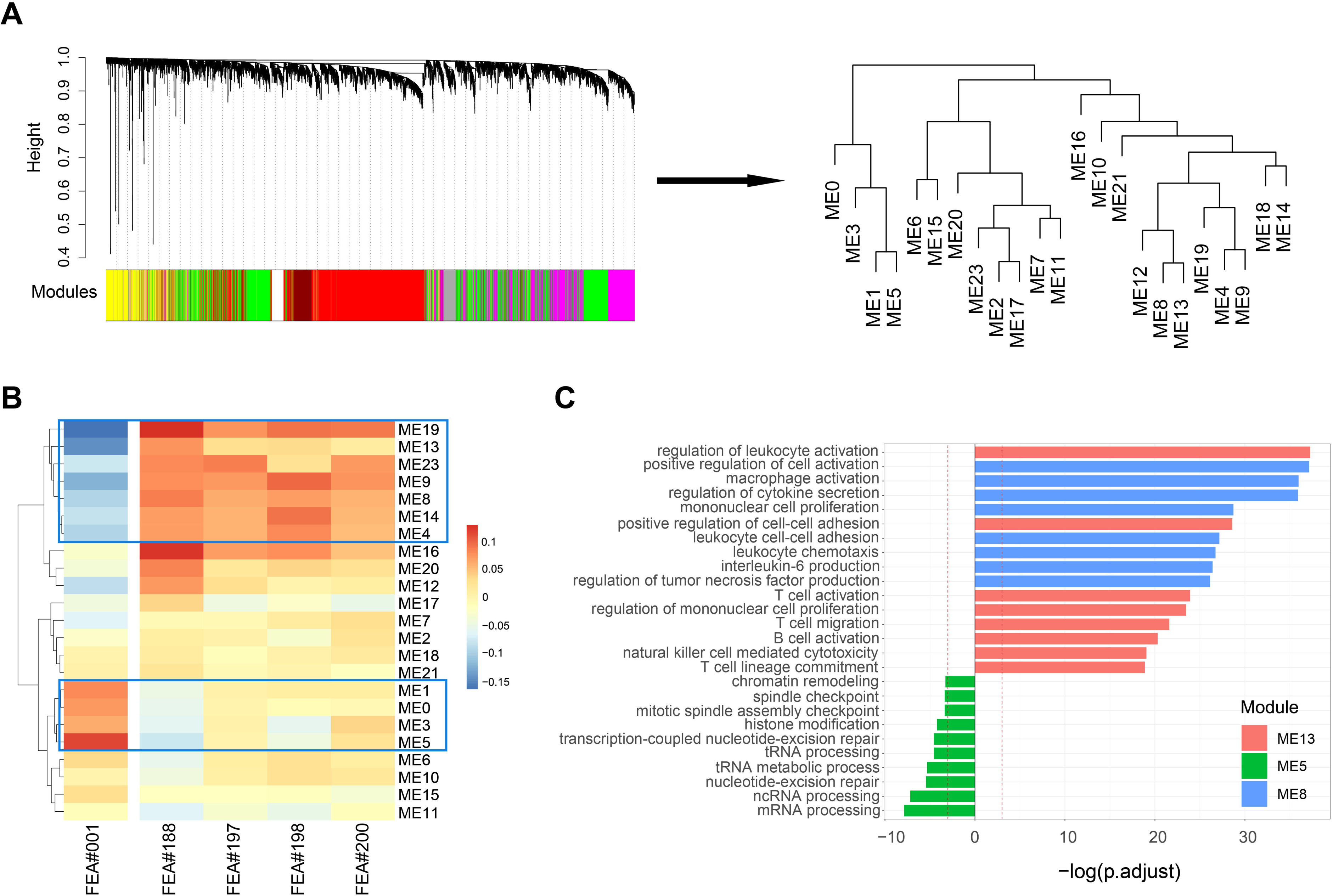
Correlation between WGCNA-identified modules and top pathological signatures. (A) Weighted gene co-expression network analysis (WGCNA) identifies gene modules with highly synergistic changes. Relationships among the modules are shown in a dendrogram. (B) Each column corresponds to one of the top five pathological signatures, and each row is one of the 24 modules identified by WGCNA. The heat map shows Spearman’s rank correlation coefficients for each pair of modules and pathological signatures, where a transition from red to blue represents positive to negative correlations. (C) Significantly-enriched Gene Ontology terms of ME5, ME8, and ME13. The dotted lines indicate the level with an adjusted *P* value of 0.05.

Spearman’s rank correlation between the 24 WGCNA modules and the top five pathological signatures showed that 11 out of 24 modules were of interest. Among them, seven modules, including ME19, ME13, ME23, ME9, ME8, ME14, and ME4, were found to be negatively correlated to FEA#001 but positively correlated to FEA#188/197/198/200. Conversely, four modules, including ME1, ME0, ME3 and ME5, were found to be positively correlated to FEA#001 but negatively correlated to FEA#188/197/198/200 (Figure 5B).

By referring to the significantly enriched GO terms of the correlated modules, we found that those enriched in ME13 and ME8 are mainly related to the biological processes of immune activation, such as T cell activation and regulation of leukocyte activation. As for ME5, which is positively correlated with FEA#001, some biological processes related to the genetic material activity are significantly enriched, including nucleotide-excision repair and transcription-coupled nucleotide-excision repair (Figure 5C). Representative GO terms enriched in other modules are shown in Figure S3.

These results further verify the findings in previous literature that MSI renders tumors immunogenic and is sensitive to immune checkpoint inhibitors[26].

## DISCUSSION

MSI testing can provide important information for clinical decision-making in a variety of cancers. However, the requirement of additional genetic or immunohistochemical tests limits its access to the general population. In this study, we developed a deep learning model which we termed Ensemble Patch Likelihood Aggregation (EPLA) to predict MSI status of CRC directly from histopathology images that are ubiquitously available in clinical practice, making it possible for every patient with a pathological diagnosis to receive an MSI evaluation. Furthermore, we propose the use of transfer learning for model fine-tuning in a different population, improving its generalizability. We also explored the model interpretability from the perspective of genome and transcriptome association, giving a molecular biological explanation of our model.

In the development of a state-of-the-art method, Kather et al. proposed a deeplearning based majority voting method to predict MSI from histology under the assumption that all patches contribute equally to the prediction of MSI status. Such assumptions may not be valid and could limit the prediction accuracy. In practice, although hundreds of patches are tiled from each WSI, most of them do not contribute much to the final prediction. In contrast, only a few key patches make the majority contribution. Our MIL-based model has the ability to automatically adjust the contribution of each patch to the overall WSI-level prediction in a learnable way by giving key patches higher weights, resulting in higher performances over the DL-based MV method in terms of AUC, sensitivity, and specificity.

In clinical practice, different data sets could be vastly different due to the disparities between patient populations and data acquisition processes, resulting in a large performance gap for AI algorithms. For example, in Kather’s study, an MSI classifier trained on TCGA, which is mainly made up of Western populations, and only achieved an AUC less than 0.70 in the KCCH cohort, a Japanese cohort[18]. Furthermore, the histology slides in TCGA-COAD are flash-frozen slides that utilize water crystallization during the freezing process, often resulting in an altered appearance of the tissue structure as compared to the FFPE slides used in Asian-CRC which provide more tissue structure clarity. This data difference could not be effectively eliminated by only color normalization, indicating that more advanced techniques, such as transfer learning, are necessary. As expected, EPLA showed a performance degradation in Asian-CRC by simply applying the model trained on TCGA-COAD, but the results improved significantly after transfer learning. It is of clinical significance that using only a minority of the new domain data for model finetuning can already improve the AUC to a satisfactory level and further improvement can be expected if even more data were included, which proves that our model can be easily generalized to the complicated clinical environment, regardless of race, preparation techniques, and data acquisition techniques.

Deep learning models are often criticized for their poor interpretability, especially in mission-critical applications, such as healthcare. Only those models with certain interpretability can be understood, verified, and trusted by clinicians in clinical practice[30]. The genomic association between DNA repair pathways and MSI cancers, which is exquisitely sensitive to ICB, has been verified previously[31]. Moreover, MSI and its resultant TMB have been reported to underlie the response to PD-1 blockade immunotherapy[32, 33], and the INDEL mutation load is particularly associated with the extent of the response[26]. Despite advances in our understanding of MSI at the genomic level, the process and mechanisms at the transcriptomic level remain relatively understudied. Previous studies have described the molecular portrait of MSI from the perspective of gene expression level and suggest that the mutation events of the MMR pathway are responsible for the under-expression of its gene members[31]. Inspired by these discoveries, we confirmed the correlation between the pathological signatures of the model and these known biomarkers of MSI. In addition, we also observed a correlation with the expression level of immune pathways, a finding which is consistent with better prognosis and a more favorable response to ICB in MSI patients.

The nature of AI models has limitations in our model. The performance of deep learning models largely depends on the size and quality of the training set. We still need to expand the training data to improve the accuracy and generalizability of the model. Although the model has been verified in TCGA and an Asian cohort respectively, a large prospective clinical trial is necessary before we can deploy it as a routine MSI testing method in clinical practice.

### Conclusions

In this study, we developed a model which we termed Ensemble Patch Likelihood Aggregation (EPLA) for MSI prediction directly from pathological images without the need for genetic or immunohistochemical tests. Using these images allows the evaluation of MSI status in many more patients than was previously possible. Through the model, we identified five pathohistological imaging signatures to predict the MSI status. The reliability of the model was verified in two independent cohorts and the interpretability of the model was illustrated by exploring the correlation between the pathological signatures and genomic and transcriptomic characterizations. Ongoing work is attempting to further validate our model in large-cohort, prospective clinical trials.

## Supporting information

Table S1. Summary of the TCGA-COAD and Asian-CRC cohorts.

Table S2. Sensitivity and specificity of different models with optimal cutoff evaluated in the TCGA-COAD test set.

Figure S1. Scale independence and mean connectivity of the WGCNA network for soft threshold determination. The recommended power value of the optimal

Figure S2. Adjacency heatmap of the WGCNA identified modules. The heat map shows Spearman’s rank correlation coefficients for each pair of modules, wh

Figure S3. Representative Enriched Gene Ontology (GO) Terms in Modules of Interest.

Table S3. GO Analysis in Module

## Acknowledgement

We would like to thank the Shanghai Tongshu Biotechnology Co., Ltd. for MSI detection.

## Author contributions

R.C., F.Y. and S.C.M. performed the experiments; Z.Y.D., J.H.Y. and L.L. designed the experiments; Y.L., H.B.Z., Y.W.L., and J.J.K. delineated the ROI of TCGA and Asian-CRC cohorts; Y.Z., W.J.L., T.X.W. and W.M.T tiled the WSIs and preprocessed the patches; Y.L. and W.J.C. collected external validation histopathology images and helped identify samples validated by MSI detection; R.C., F.Y., W.J.H. and S.C.M. contributed to the analysis of the data; Z.Y.D., J.H.Y. and L.L. conceived and directed the project; Z.Y.D., S.C.M., J.H.Y., L.L., Y.Z. and F.Y. wrote the manuscript with the assistance and feedback of all the other co-authors.

## Ethics approval

This study was approved by the Institutional Ethical Review Boards of Nanfang Hospital

