## Supplementary material for "A transferrable and interpretable multiple instance learning model for microsatellite instability prediction based on histopathology images": Table S1. Summary of the TCGA-COAD and Asian-CRC cohorts.

| **Cohort** | **Material** | **Annotated WSI** | **MSS** | **MSI** | **Patches** |
| --- | --- | --- | --- | --- | --- |
|  |  |  |  |  | **min, 25%, 50%, 75%, max** |
| TCGA-COAD | Frozen slides | 429 | 358 | 71 | 22, 143, 229, 398, 2357 |
| Asian-CRC | FFPE | 785 | 621 | 164 | 5, 179, 338, 608, 3718 |

Abbreviations: FFPE: formalin-fixed paraffin-embedded; WSI: whole slide image; MSI: microsatellite instability; MSS: microsatellite stability.
