## Supplementary material for "A transferrable and interpretable multiple instance learning model for microsatellite instability prediction based on histopathology images": Table S2. Sensitivity and specificity of different models with optimal cutoff evaluated in the TCGA-COAD test set.

|  | **DL-based MV** | **PALHI pipeline** | **BoW pipeline** | **EPLA** |
| --- | --- | --- | --- | --- |
| optimal cutoff | 0.57 | 0.63 | 0.62 | 0.68 |
| Sensitivity | 0.82 | 0.86 | 0.73 | 0.91 |
| Specificity | 0.75 | 0.76 | 0.9 | 0.77 |

Abbreviations: DL-based MV: deep-learning based majority voting; PALHI: PAtch Likelihood Histogram; Bag of Words (BoW); EPLA: Ensembled Patch Likelihood Aggregation.
