## Supplementary figures and images for "A transferrable and interpretable multiple instance learning model for microsatellite instability prediction based on histopathology images"

### Figure S1. Scale independence and mean connectivity of the WGCNA network for soft threshold determination. The recommended power value of the optimal

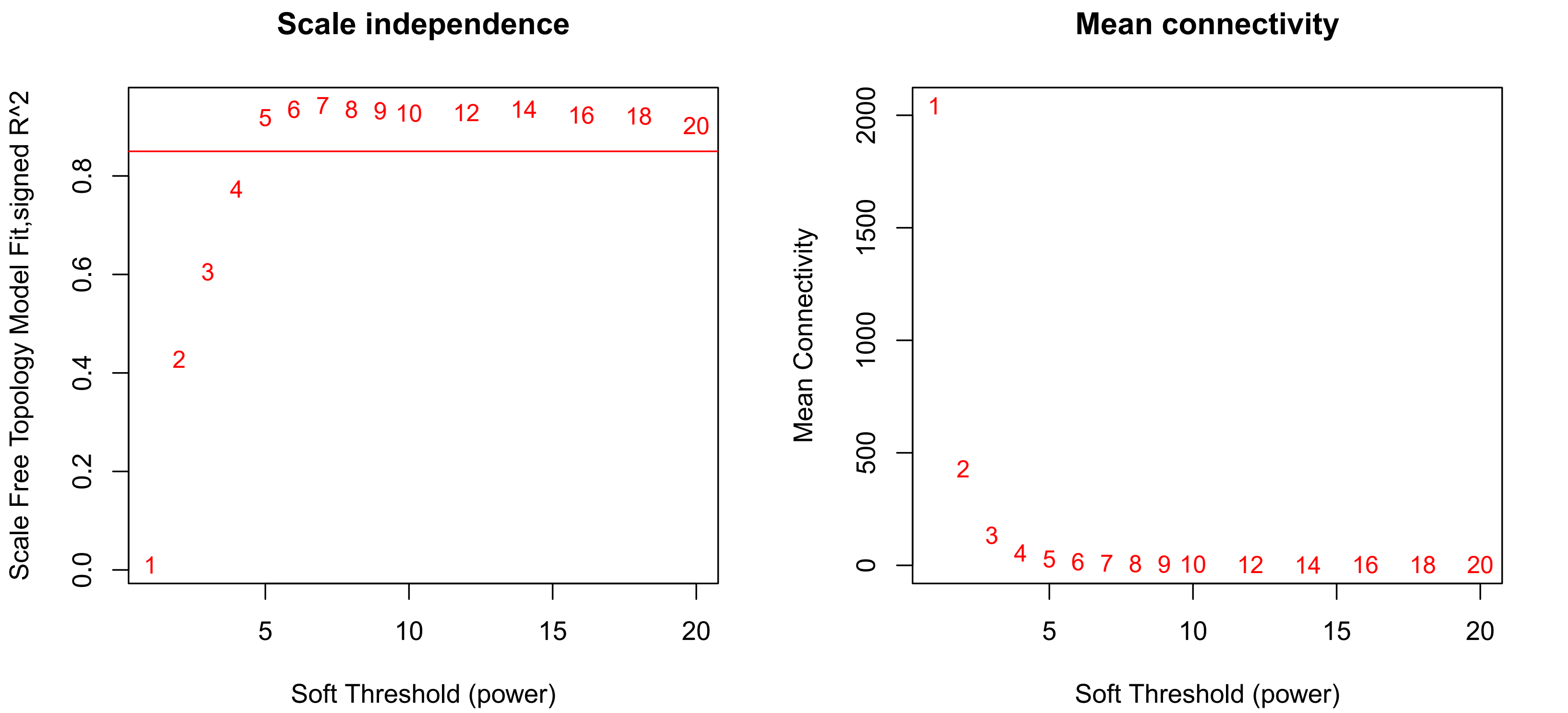

### Figure S2. Adjacency heatmap of the WGCNA identified modules. The heat map shows Spearman’s rank correlation coefficients for each pair of modules, wh

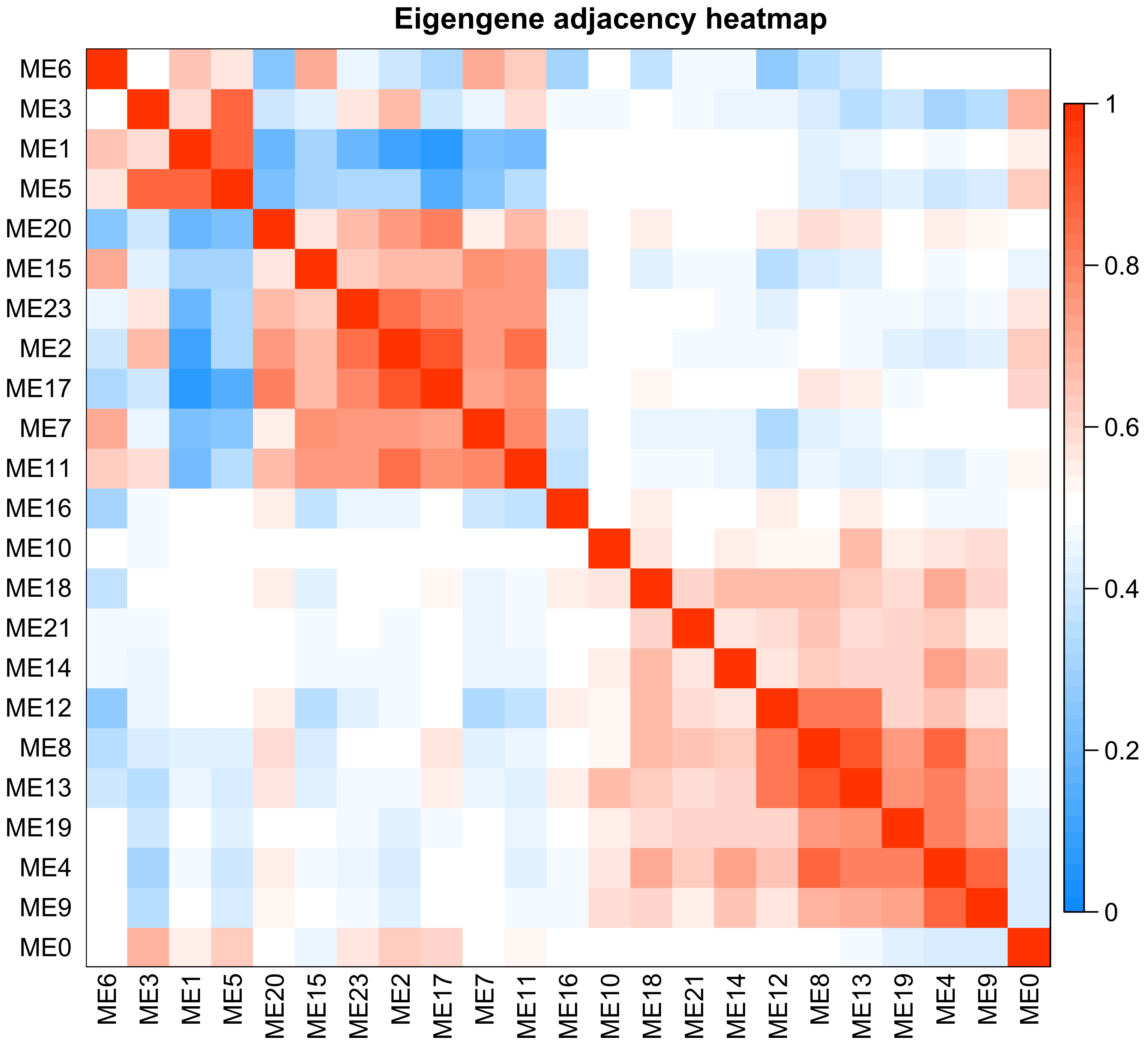

### Figure S3. Representative Enriched Gene Ontology (GO) Terms in Modules of Interest.

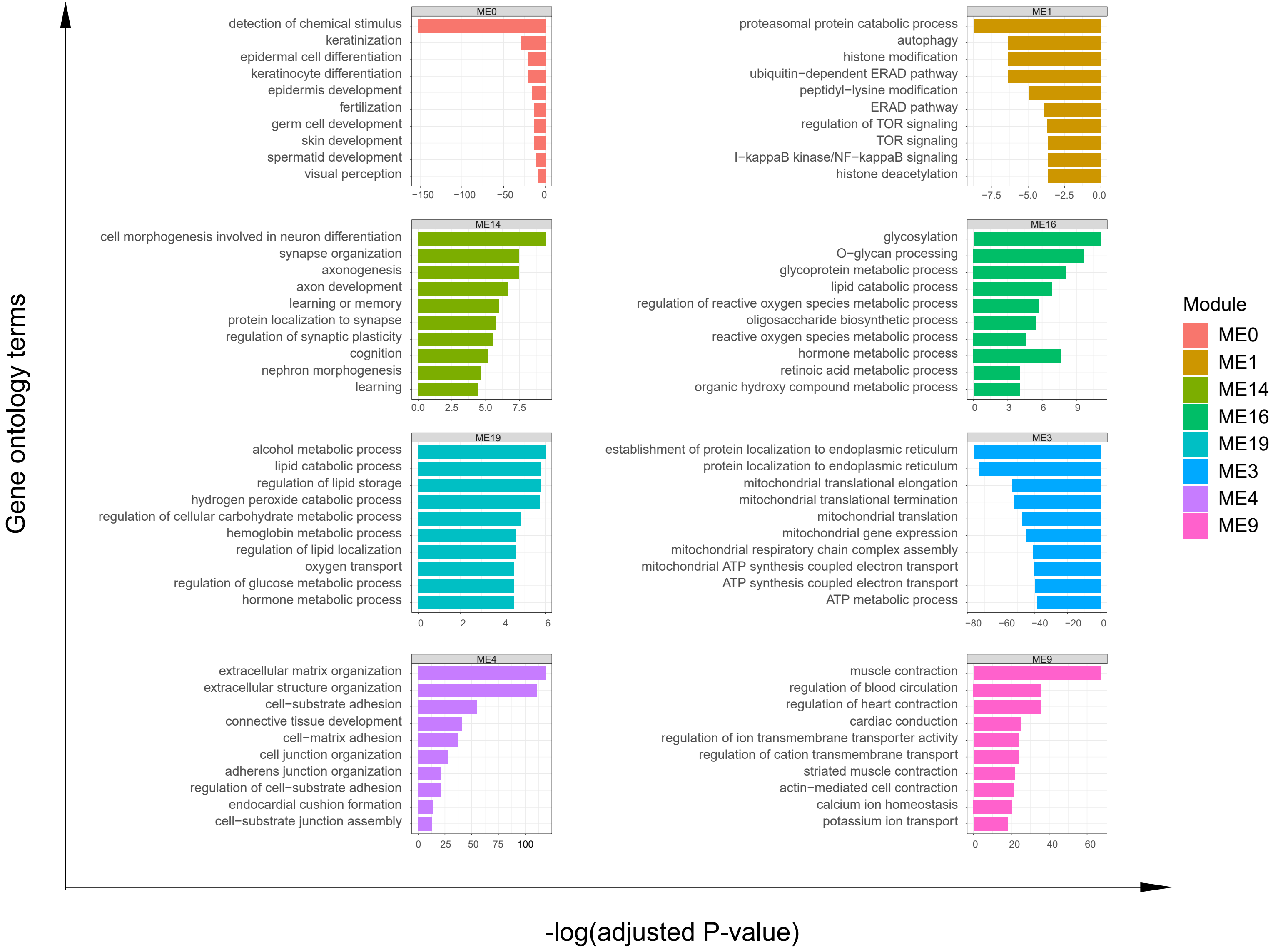
